# Spinal reconsolidation engages non-ionotropic NMDA receptor signaling to reverse pain hypersensitivity

**DOI:** 10.1101/2021.09.08.458120

**Authors:** Abigail J. D’Souza, David Rodriguez-Hernandez, Hantao Zhang, David He, Maham Zain, Samuel Fung, Laura A. Bennett, Robert P. Bonin

**Author notes:** Corresponding Author:* Robert P. Bonin, PhD., Leslie Dan Faculty of Pharmacy, University of Toronto, Toronto, Ontario, Canada. These authors contributed equally (Abigail J. D’Souza, David Rodriguez-Hernandez, Hantao Zhang). **Contributions** All authors contributed to the project design, conducted experiments and analyzed data. AJD, DRH, DH performed and analyzed electrophysiology; HZ collected and analyzed western blot data; HZ, MZ, SF, LA, and RPB performed and analyzed behavioural experiments. AJD, DRH, HZ, and RPB wrote the manuscript and all authors edited and approved the final manuscript. **Disclosure** AV-101 was provided by VistaGen Therapeutics. VistaGen Therapeutics did not provide input or influence on the design, analysis, or reporting of results.

## Abstract

Reconsolidation enables the activity-dependent modification of memory traces and has been used to reverse addiction, fear memory, and pain hypersensitivity in animal models. We demonstrate that non-ionotropic NMDA receptor signalling in the spinal dorsal horn is sufficient to reverse pain hypersensitivity and necessary for pain modulation by spinal reconsolidation. These findings reveal a key process by which reconsolidation disrupts memory traces that may be exploited in the treatment of pain and other disorders.

## Main Body of Manuscript

Many treatments for pain, such as opioids, dampen neuronal activity to suppress nociceptive processing but do not address underlying causes of pain. Central sensitization or nociplastic pain represents a particularly intractable source of chronic pain, as this “memory trace”^1,2^ of pain can stably persist long after being triggered by an insult or injury^3^. Previous work has demonstrated that spinal sensitization and hyperalgesia can be reversed in a process analogous to memory reconsolidation: a protein synthesis-dependent process by which memory traces can be updated or modified^4,5^. Preventing protein synthesis after reactivation of sensitized nociceptive networks resulted in the reversal of dorsal horn long-term potentiation and mechanical hyperalgesia^4^. This finding suggests that exploiting the mechanisms of reconsolidation may be an effective approach to treat central changes contributing to pathological pain.

Reconsolidation can be characterized as having distinct processes of protein synthesis and protein degradation that underlie synaptic potentiation and depotentiation, respectively^6^. There has been considerable research into the molecular conditions necessary for the induction of protein synthesis and restabilization of memory traces in reconsolidation^6-8^. However, it is unknown which signalling cascades contribute to the degradation of memory traces that is enabled by reconsolidation and of therapeutic relevance. We previously demonstrated that reconsolidation in spinal nociceptive networks requires activation of NMDA receptors^4^. Recent work has indicated that NMDA receptors can signal in a non-canonical, non-ionotropic manner to reduce synaptic efficacy^9-13^. We examined whether non-ionotropic NMDA (NI-NMDA) signalling is sufficient to reverse sensitization and hyperalgesia, and whether it contributes to the depotentiation process of spinal reconsolidation.

To test whether NI-NMDA can reverse hyperalgesia, mechanical sensitization was induced by intraplantar injection of capsaicin in mice. Three hours after capsaicin injection, NI-NMDA was induced by intrathecal co-injection of the NMDA (75 μM) and the NMDA glycine site antagonist, 7-chlorokynurenic acid (7-CK, 250 μM), which reduced sensitization 3 hours after treatment as indicated by an increase paw withdrawal threshold (PWT; Figure 1A). Intrathecal administration of NMDA or 7-CK alone did not affect PWT. The reversal of hyperalgesia by intrathecal 7-CK was observed in both male and female mice (Supplemental Figure 1A). Similarly to previous work, in which pain reconsolidation was previous induced by a second intraplantar capsaicin injection^4^, we also observed a reduction in PWT when a second capsaicin injection was preceded by intrathecal injection of 7-CK, another potent glycine site antagonist L-689,560 (50 μM), or intraperitoneal injection of the blood-brain barrier permeable prodrug of 7-CK, AV-101^14^ (400 mg/kg; Fig 1B,C). In contrast, intrathecal administration of the NMDA glutamate binding site antagonist APV (250 μM) alone or co-administered with 7-CK prior to the second capsaicin injection did not affect PWT (Fig 1D), indicating that the reversal of hyperalgesia is not caused by general inhibition of the NMDA receptor. Additionally, this effect was independent from GluK1 kainate receptors or metabotropic glutamate receptor mGluR5, which have been previously shown to modulate spinal plasticity^15,16^ (Supplemental figure 1B). The degree to which NI-NMDA activity reversed hyperalgesia varied with the concentration of intraplantar capsaicin, indicating that spinal NI-NMDA signalling is driven by nociceptor activity (Fig 1E).

**Figure 1.**
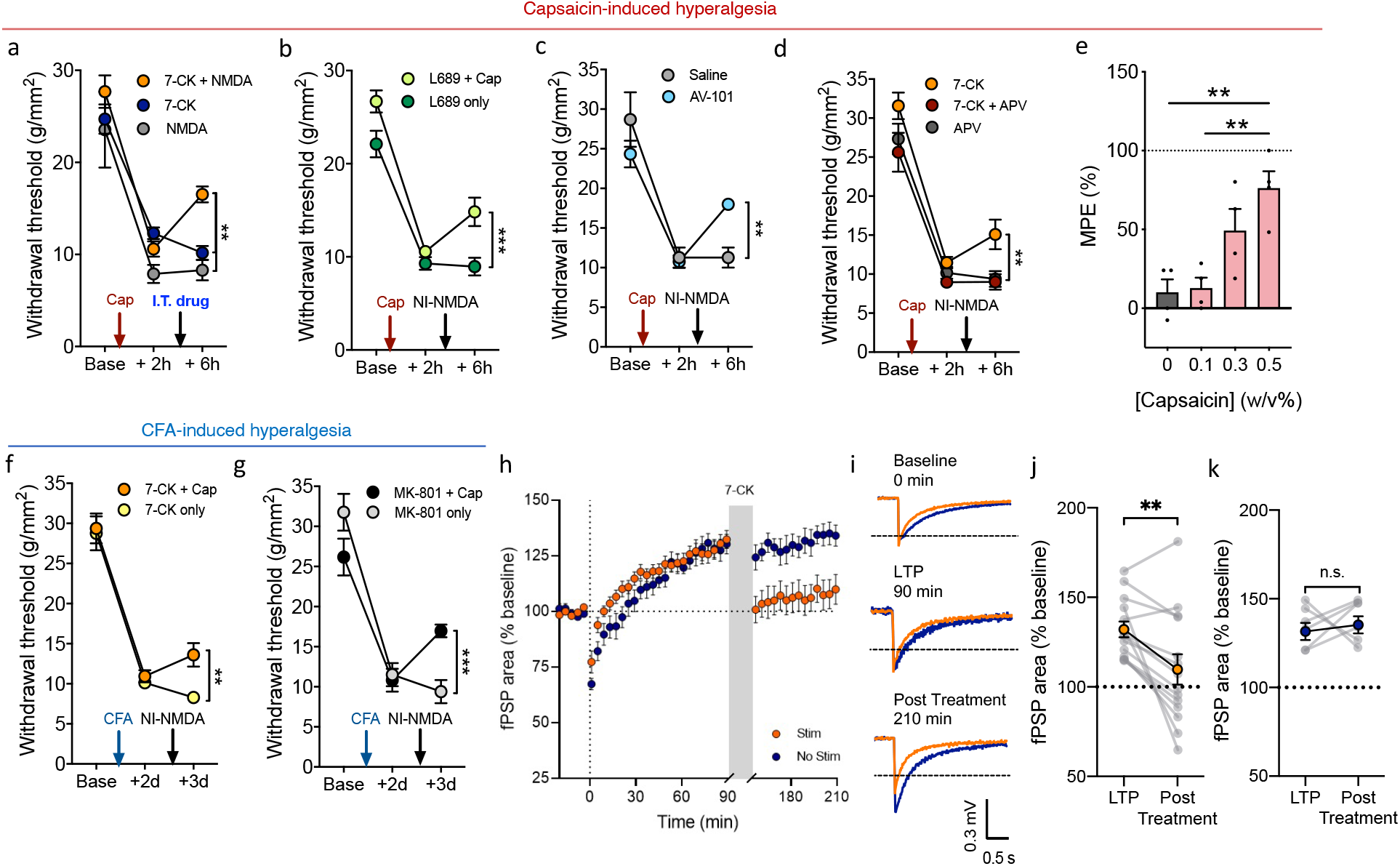
Non-ionotropic NMDA receptor signaling renders both mechanical hyperalgesia and LTP labile and reversible. a) Mechanical withdrawal thresholds were assessed in mice at baseline (Base) and after injection of capsaicin (‘Cap’, red arrows). NI NMDA activation (i.t. 7-CK + NMDA) partially reversed sensitization caused by capsaicin, while i.t. 7-CK or NMDA alone did not. n = 12 per group, ** P < 0.01. b) NI-NMDA can also be induced by potent glycine site antagonist, L-689,560 (i.t.; L689) and a 2^nd^ intraplantar capsaicin injection to cause the reversal of hyperalgesia. n=8 per group, ***P < 0.001 c) Similar to 7-CK and L689, AV-101 caused reversal of hyperalgesia when combined with a second capsaicin injection, but not when given without capsaicin injection. n = 5, ** P <0.01 d) NI-NMDA induced by i.t. 7-CK and a 2nd intraplantar capsaicin injection was prevented by co-administration of 7-CK and APV (i.t). n = 6, ** P <0.01 e) Antihyperalgesic effect mediated by AV-101 at different concentrations of 2^nd^ intraplantar capsaicin injection (0; 0.1; 0.3; 0.5 w/v%) expressed as percentage of maximum possible effect (MPE) n = 4 per group, ** P <0.01. Two days after the induction of hyperalgesia by plantar CFA, i.t. injection of (f) 7-CK n = 6 per group, ** P <0.01 or (g) MK-801 n=6, ***P <0.001, 30 min prior to a plantar injection of capsaicin caused a significant reduction of hyperalgesia when mechanical sensitivity was tested the following day. h) LTP of fPSPs induced by 2-Hz stimulation at time = 0 min, followed by 7-CK administration from 90 min to 150 min (shaded area) with (orange circles) or without (blue circles) electrical stimulation of dorsal roots. i) Representative traces of fPSPs recorded during baseline (0 min; top), at 90 min (middle) and 210 min (bottom) post 2-Hz stimulation. j-k) The magnitude of fPSP potentiation was compared before (LTP) and after (Post Treatment) 7-CK administration. Stim: n = 14, t _(13)_ = 3.514, ** *p* = 0.0038; No Stim: n = 6, t _(5)_ = 0.4343, *p* = 0.6822.

To test whether spinal NI-NMDA can produce long-lasting analgesic effects, we further used intraplantar injection of complete Freund’s adjuvant (CFA) that induces inflammatory hyperalgesia lasting up to two weeks. Two days after injection of CFA, NI-NMDA was induced by intrathecal injection of 7-CK or the NMDA receptor pore-blocker MK-801 (2 mM) followed by intraplantar injection of capsaicin. In both cases, hyperalgesia was reduced when assessed 24-hours after treatment (Fig 1E,F), indicating a long-lasting antihyperalgesic effect of NI-NMDA.

Hyperalgesia is associated with long-term potentiation (LTP) in the spinal dorsal horn^17^. Because the induction of NI-NMDA can lead to a reduction in synaptic efficacy and synaptic loss^9,12^, we next tested whether NI-NMDA can reverse LTP in the spinal dorsal horn. After the induction of LTP in spinal cord explants from adult male mice, 7-CK was applied to the bath (100 μM) while dorsal roots were electrically stimulated, leading to a reduction in evoked field post-synaptic potentials (fPSPs; Fig 2A-C). Interestingly, bath application of 7-CK with or without stimulation did not affect fPSPs in the absence of prior LTP (Supplemental Figure 2), indicating that NI-NMDA causes depotentiation rather than a non-specific reduction in synaptic strength or long-term depression. Additionally, blockade of NMDA receptors with APV (100 μM) did not cause an activity-dependent reversal of LTP (Supplemental Figure 3A), while the induction of NI-NMDA with L-689,560 (50 μM) also reduced LTP (Supplemental Figure 3B). The reduction of LTP by NI-NMDA was observed in spinal cord explants from both male and female mice (Supplemental Figure 4A) as well explants from both C57BL/6N and CD-1 mice (Supplemental Figure 4B), suggesting NI-NMDA is a conserved mechanism of depotentiation in the dorsal horn.

**Figure 2.**
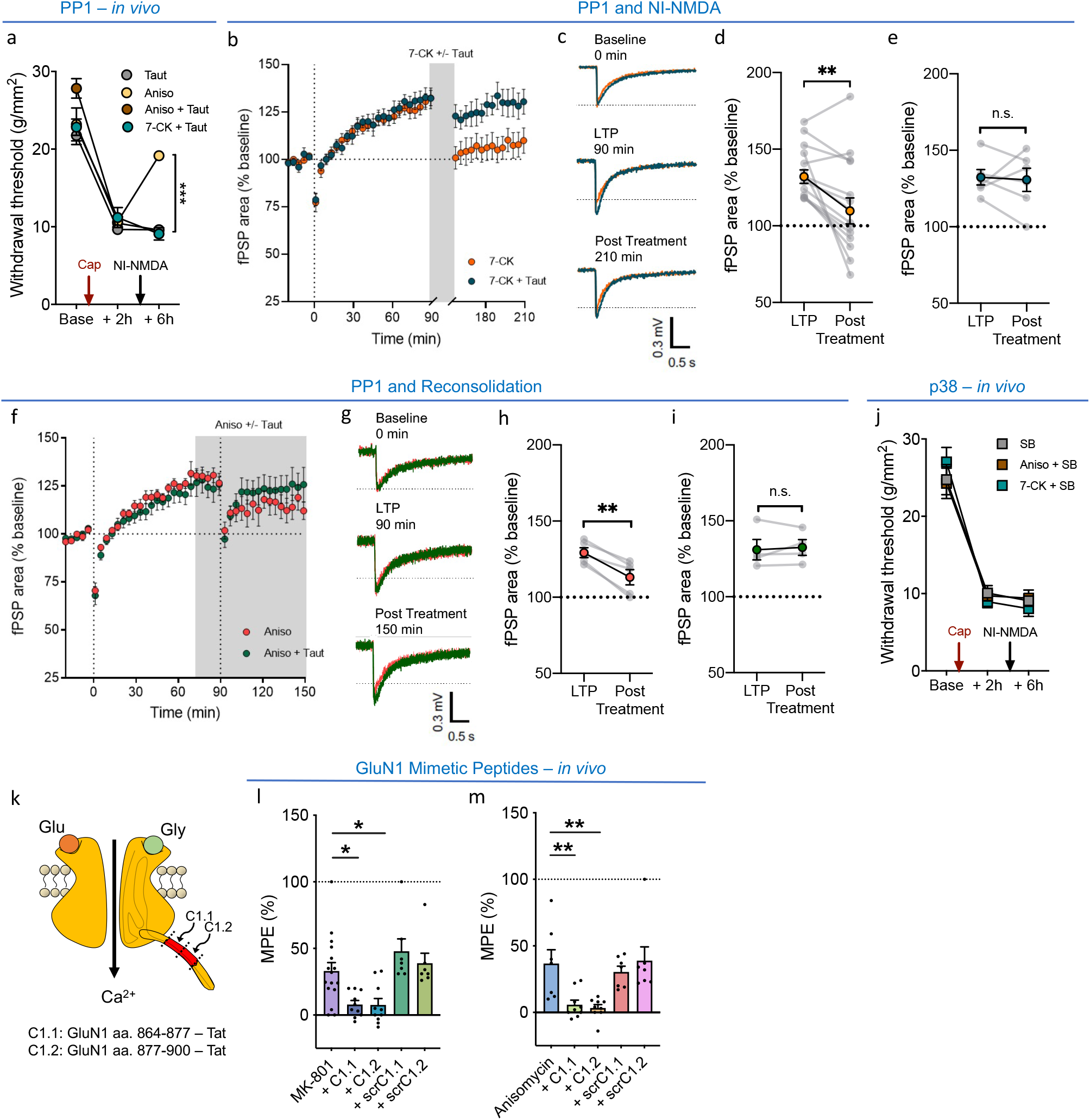
Reversal of hyperalgesia and dorsal horn LTP induced by NI-NMDA and reconsolidation require overlapping mechanisms downstream of NMDA receptor activation. a) Intrathecal tautomycetin (Taut) administered with 7-CK or anisomycin (Aniso) given prior to a second capsaicin injection (black arrow) prevents reversal hyperalgesia n=6 per group, ***P < 0.001 b) LTP of fPSPs induced by 2-Hz stimulation at time = 0 min, followed by 7-CK administration from 90 min to 150 min (shaded area) in the presence (cyan circles) or absence (orange circles) of tautomycetin; dorsal roots were electrically stimulated during drug administration. c) Representative traces of fPSPs recorded during baseline (top), at 90 min (middle) and 210 min (bottom) post 2-Hz stimulation. d-e) The magnitude of fPSP potentiation was compared before (LTP) and after (Post Treatment) drug administration. 7-CK: n = 14, t_(13)_ = 3.514, ** *p* = 0.0038; 7-CK + Tautomycetin: n = 6, t_(5)_ = 0.2461, p = 0.8154.f) LTP induction was followed by anisomycin administration at time = 75 min, in the presence (green circles) or absence (orange circles) of tautomycetin, and a second round of 2-Hz stimulation at 90 min. g) Representative traces of fPSPs recorded during baseline (top), at 90 min (middle) and 150 min (bottom). h-i) The magnitude of fPSP potentiation was compared before (LTP) and after (Post Treatment) the second 2-Hz stimulation. Anisomycin: n = 5, t _(4)_ = 6.155, ** *p* = 0.0035; Anisomycin + Tautomycetin: n = 4, t_(3)_ = 0.4759, *p* = 0.6667. j) Intrathecal SB-203580 (SB) administered with 7-CK or anisomycin (Aniso) given prior to a second capsaicin injection (black arrow) prevents reversal hyperalgesia n=8 per group, ***P < 0.001. k) Schematic diagram shows GluN1 C-terminal tail, it contains a C1 domain that are composed of two segments of amino acid, C1.1 and C1.2. l) Antihyperalgesic effect mediated by intrathecal MK-801 was prevented by both mimetic C1.1 and C1.2 peptides, but not by their own scrambled peptides. (MK-801: n = 16; + C1.1: n = 10, * *p* < 0.05; + C1.2: n = 10, * *p* < 0.05) m) Antihyperalgesic effect mediated by intrathecal anisomycin was prevented by both mimetic C1.1 and C1.2 peptides, but not by their own scrambled peptides. (Anisomycin: n = 7; + C1.1: n = 9, ** *p* < 0.01; + C1.2: n = 9, ** *p* < 0.01).

These findings indicate that NI-NMDA can reverse hyperalgesia and dorsal horn plasticity associated with pathological pain, making it a candidate mechanism for the activity-dependent reversal of hyperalgesia that can be induced by spinal reconsolidation. We therefore examined if protein phosphatase 1 (PP1) and p38 MAPK activity that have been associated with NI-NMDA similarly contribute to spinal reconsolidation^18^. Inhibition of protein phosphatase 1 (PP1) with tautomycetin (5 μM) prevented the reversal of hyperalgesia by both NI-NMDA and spinal reconsolidation blockade with anisomycin (47 mM; Fig 2A). We further examined the role of PP1 in the reversal of dorsal horn LTP. The reversal of LTP by NI-NMDA induced via 7-CK was prevented by bath application of tautomycetin (100 nM; Fig 2B-E). LTP was also reversed by blocking reconsolidation via bath application of anisomycin (100 μM) during stimulation, and this effect was also prevented by tautomycetin (Fig 2F-I). Similarly, inhibition of p38 MAPK with SB-203580 (100 μM) also prevented reversal of hyperalgesia by both NI-NMDA or blockade of reconsolidation induced by7-CK or anisomycin, respectively (Fig 2J). Overall, these data indicate strong parallels in the mechanisms by which NI-NMDA and reconsolidation can reverse hyperalgesia and synaptic potentiation.

We next sought to selectively disrupt NI-NMDA signalling during reconsolidation. The intracellular C-terminal tail of the GluN1 subunit is an essential hub for coordinating events downstream of NI-NMDA activity, and intracellular delivery of antibodies targeting the C-terminal tail prevents NI-NMDA signalling^11^. We therefore developed membrane-permeable peptides that mimic segments of the GluN1 C-terminal tail to interfere with NI-NMDA signalling (Figure 2K). These peptides were co-administered intrathecally (500 μM) with MK-801 (2 mM) or anisomycin prior to plantar capsaicin injection in CFA-treated mice to assess if they to block NI-NMDA or reconsolidation, respectively. The mimetic peptides C1.1 and C1.2 both prevented reversal of hyperalgesia by NI-NMDA induced with MK-801; however, membrane permeable scrambled peptides were ineffective (Figure 2L). Similarly, the mimetic peptides but not the scrambled controls also prevented reversal of hyperalgesia by spinal reconsolidation blockade, indicating that NI-NMDA signalling initiated by the GluN1 C-terminal tail is necessary for reconsolidation.

Reconsolidation requires ubiquitin-dependent degradation of synaptic proteins^19^, yet it is unknown whether similar patterns of synaptic protein degradation is induced by NI-NMDA signalling. We therefore sought to determine whether reversal of hyperalgesia by either spinal reconsolidation or NI-NMDA requires protein degradation to further establish the overlap across these processes. We found that intrathecal administration of the E1-ligase inhibitor PYR-41 (135 μM) prevented reversal of hyperalgesia induced by both processes (Fig 3). When we further examined synaptic protein degradation after induction of spinal NI-NMDA *in vivo*, we found a pattern of altered protein expression that was strikingly similar to that seen in spinal reconsolidation. Following both interventions, we observed reduced expression of AMPA receptor subunits GluA1, GluA2, SHANK3 and GKAP (Fig 3), which have been previously shown to be degraded following reconsolidation blockade^20^. However, we did not observed changes in PSD-95 (Fig 3), CaMKII expression or its phosphorylation form (Supplemental Fig 5a & 5b).

**Figure 3.**
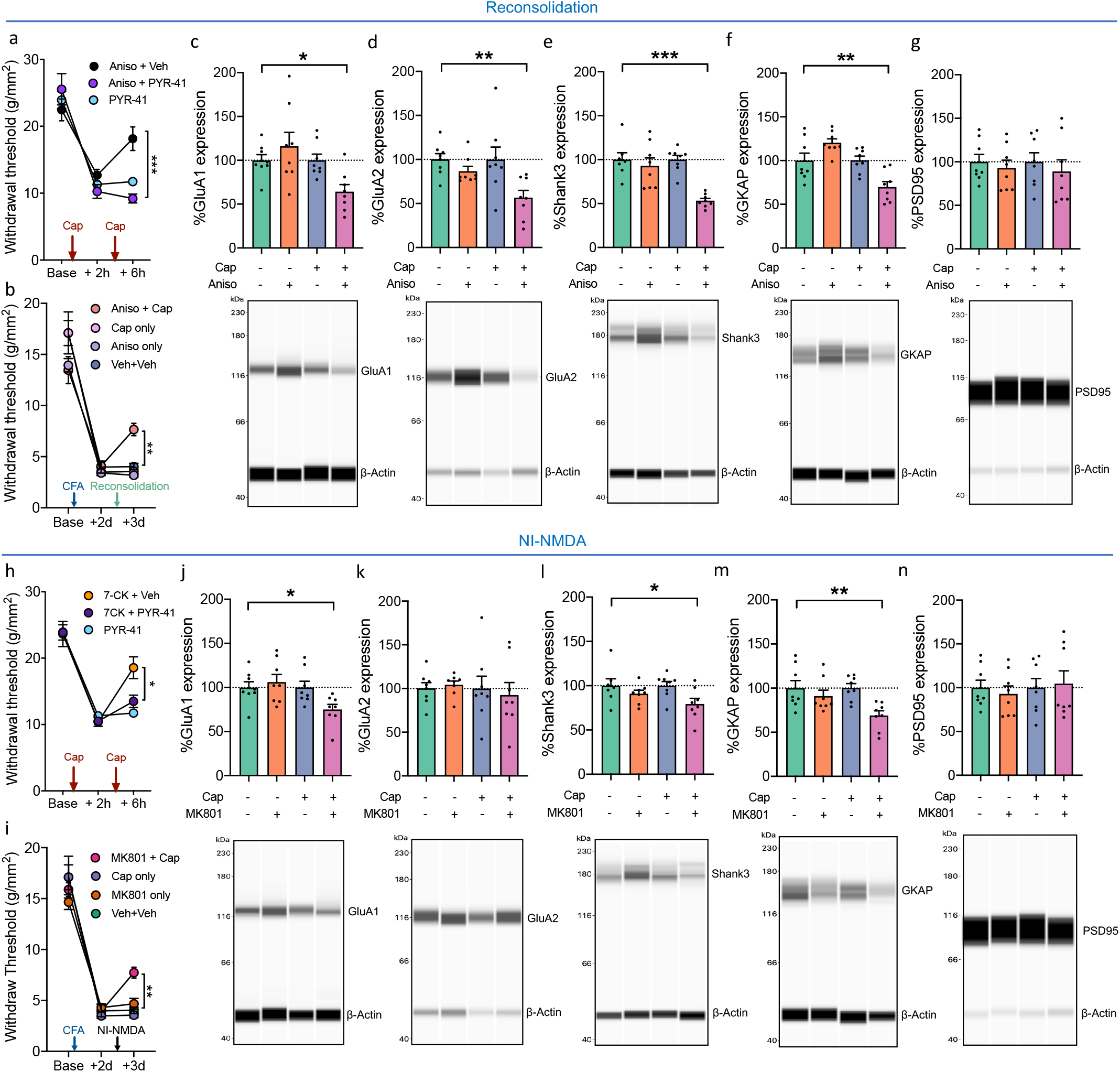
Protein ubiquitination and degradation underlies reversal of hyperalgesia by spinal reconsolidation and NI-NMDA. a) Intrathecal administration of PYR-41 (E1 ubiquitin ligase inhibitor) with anisomycin (Aniso) prior to a second capsaicin injection prevents the reversal of hyperalgesia. (n = 6 per group, *** *p* < 0.001). b) Intrathecal administration of anisomycin prior to plantar capsaicin injection is associated with reversal of hyperalgesia (n = 8 per group, ** *p* < 0.01). Spinal cords from mice in (b) were used to generate data in (c – g). Reversal of hyperalgesia by spinal reconsolidation was associated with a reduction in synaptosome protein levels of (c) GluA1 (n = 8 per group, * *p* < 0.05), (d) GluA2 (n = 8 per group, ** *p* < 0.01), (e) Shank3 (n = 8 per group, *** *p* < 0.001), (f) GKAP (n = 8 per group, ** *p* < 0.01), but no change in (g) PSD-95 (n = 8 per group). (h) Similarly, intrathecal administration of PYR-41 (E1 ubiquitin ligase inhibitor) with 7-CK prior to a second capsaicin injection prevents the reversal of hyperalgesia. (n = 6 per group, * *p* < 0.05). i) Intrathecal administration of 7-CK prior to plantar capsaicin injection is associated with reversal of hyperalgesia (n = 8 per group, \*\**p* < 0.01). Spinal cords from mice in (i) were used to generate data in (j – n). Reversal of hyperalgesia by NI-NMDA was similarly associated with a reduction in synaptosome protein levels of (j) GluA1 (n = 8 per group, * *p* < 0.05), (l) Shank3 (n = 8 per group, * *p* < 0.05), (m) GKAP(n = 8 per group, ** *p* < 0.01), but no change was observed in (k) GluA2 (n = 8 per group) or (g) PSD-95 (n = 8 per group).

Taken together, these results indicate that NI-NMDA is a conserved mechanism across different strains and sex of mice that induces synaptic depotentiation to reverse hyperalgesia. Crucially, the necessity of NI-NMDA activation in reconsolidation indicates that NI-NMDA signalling is responsible for the erasure of memory traces enabled during reconsolidation. These data therefore link both NMDA receptor activity to both the synthesis and degradation of synaptic proteins in reconsolidation^6,7^, making the NMDA receptor a master signalling hub of reconsolidation. It is notable that NI-NMDA had no effect on synaptic strength in the absence of LTP, suggesting even widespread activation of this depotentiation mechanism in the dorsal horn may have negligible impact on non-potentiated sensory processing networks. We accordingly propose that engaging this conserved mechanism of depotentiation may provide a novel therapeutic strategy for directly and selectively targeting maladaptive plasticity associated with chronic pain.

## Materials and Methods

### Animal Studies

All behavioral experiments were conducted in accordance with the guidelines established by the Canadian Council for Animal Care and Local Animal Care Committee. Adult (>12 weeks old) male C57BL/6N mice were used for experiments except where indicated otherwise. Mice were kept on a 14:10 light:dark cycle in groups of 1 to 4 mice per cage with food and water provided *ad libitum*. All experiments were conducted on naïve mice and started prior to 10:00. Mechanosensitivity was measured using the SUDO up-down method with von Frey hairs to estimate the 50% withdrawal threshold in pressure units (g·mm^−2^)^21^. Care was taken to avoid the injection site when testing mechanosensitivity. Mechanical hyperalgesia was induced by intraplantar injection of capsaicin (5 μl, 0.5% w/v). Inflammatory hyperalgesia was induced by intraplantar injection of Complete Freud’s Adjuvant (10 ul) and all intraplantar and intrathecal injections (5 μl) were performed under light (< 3 min) isoflurane anesthesia as previously described^22^. In behavioural experiments, mice were excluded if they did not exhibit a reduction in withdrawal threshold greater than 10% after sensitization. Animals in which intrathecal injection did not produce an obvious tail flick were excluded from analysis. Animals were randomly assigned to treatment groups and the experimenter was blinded during testing and data analysis. Behavioral data were analyzed as percentage of maximum possible effect (MPE) using the formula 100%·(test 3 – test 1)·(baseline – test 1)^−1^.

### Electrophysiology

Electrophysiological recordings of dorsal root-evoked post-synaptic field potentials (fPSPs) were made using a whole spinal cord tissue preparation. Adult male C57Bl/6N mice (Charles River, Sherbrooke, Quebec, Canada) were anesthetized with chloral hydrate (400 mg / kg) and perfused with ice-cold sucrose substituted artificial cerebrospinal fluid (sucrose aCSF; 50 mM sucrose, 92 mM NaCl, 15 mM D-glucose, 26 mM NaHCO_3_, 5 mM KCl, 1.25 mM NaH_2_PO_4_, 0.5 mM CaCl_2_, 7 mM MgCl_2_ and 1 mM kynurenic acid, bubbled with 95%/5% oxygen/CO_2_). In some experiments spinal explants were isolated from adult male CD-1 mice or adult female C57BL/6N mice (Charles River). The lumbar spinal segment was removed and immersed in ice-cold sucrose aCSF, after which nervous tissue was quickly isolated via laminectomy. Ventral roots and connective tissue were removed from the spinal cord and the tissue was placed in room temperature aCSF (124 mM NaCl, 10 mM D-glucose, 26 mM NaHCO_3_, 3 mM KCl, 1.25 mM NaH_2_PO_4_, 2.6 mM CaCl_2_ and 1.3 mM MgCl_2_, bubbled with 95% oxygen:5% CO_2_) for 1 hr before experimentation. During experiments, the tissue was perfused with aCSF at room temperature at a flow rate of 6-8 ml min^-1^.

fPSPs were recorded via a borosilicate glass electrode inserted into the dorsal side of the spinal cord at the dorsal root entry zone. Electrodes were inserted superficially to a depth of no more than 125 μm from the dorsal surface of the spinal cord, measured with an MPC-200 manipulator (Sutter Instrument Company, Novato, CA, USA). Electrodes had a tip resistance of 4-5 MOhm when filled with aCSF. fPSPs were evoked by electrical stimulation of the dorsal root using a suction electrode that is pulled from borosilicate glass, filled with aCSF, and placed near the cut end of the dorsal root. Field potentials were amplified with a Multiclamp 700A amplifier (Molecular Devices, Sunnyvale, CA, USA), digitized with a Digidata 1322A digitizer (Molecular Devices), and recorded using pClamp 10 software (Molecular Devices). Data were filtered during acquisition with a low pass filter set at 2 kHz and sampled at 10 kHz.

Test stimuli were presented every 30 sec to evoke fPSPs. The stimulus intensity was sufficient to activate C-fibers as indicated by the appearance of a third distinct fiber volley after the stimulus artifact, while a slightly (20%) higher intensity was used to induce long-term potentiation (LTP). LTP was induced by low frequency stimulation (2 Hz, 2 min) of the dorsal root as described previously^4^. This LTP likely reflects an increase in the net postsynaptic activation of superficial dorsal horn neurons since supraspinally projecting neurons only comprise a small percentage of the neurons in this area^23^. After a stable baseline recording (20 min), the LTP protocol was presented at time = 0 min. Experimental drugs were added to the aCSF at 90 min and washed out at 150 min, for a total application of 1 hr. For reconsolidation experiments, anisomycin and tautomycetin were added to the aCSF at 75 min, and a second round of 2-Hz stimulation was presented at time = 90 min. In experiments where LTP was not induced, 7-CK was administered at 0 min and washed out at 60 min. Data were analyzed using ClampFit 10 software (Molecular Devices). The area of fPSPs relative to baseline was measured from 0−800 ms after the onset of the fPSP. All drugs and chemicals for electrophysiology solutions were purchased from Millipore-Sigma Canada (Oakville, ON, Canada).

### Isolation of synaptic fractions

For biochemical studies, adult C57Bl/6N mice were anesthetized with intraperitoneal (i.p.) injection of 400 mg/kg chloral hydrate (Sigma) and transcardiac perfused with ice-cold, sucrose-substituted aCSF (50 mM sucrose, 92 mM NaCl, 15 mM D-glucose, 26 mM NaHCO_3_, 5 mM KCl, 1.25 mM NaH_2_PO_4_, 0.5 mM CaCl_2_, 7 mM MgCl_2_ and 1 mM kynurenic acid, bubbled with 95% oxygen:5% CO_2_). The spinal cord was removed from the lumbar spinal column via laminectomy under aCSF and immediately placed the cord in ice-cold sucrose aCSF.

The lower lumbar region of the spinal cord was placed on a bed of dry ice/metal plate and allowed to freeze after which it was cut along the frontal plane to separate the dorsal horn section. Isolation of the synaptic fraction was performed as previously described, with minor modifications^24^. Briefly, tissues were homogenized in microtube homogenizer (Bel-art™) in 300 ul lysis buffer (0.32M sucrose, 5mM HEPES, pH 7.4) supplemented with one tablet of Roche Complete Mini-Protease Inhibitor (Millipore Sigma) and 100 μl Phosphatase Inhibitor Cocktail 2 & 3 (Millipore Sigma). Aliquots of lysates were saved for further analyses as total homogenates. The rest of lysates went through two spins at 4 °C (10 min at 1,000 x *g* and 20 min at 12,000 x *g*) to obtain crude synaptic fractions. Supernatant containing the light membrane fraction and soluble enzymes (S2) was immediately frozen for further analyses as non-synaptic fraction, and pellet containing crude synaptosome (P2) was resuspended in 200 ul of 0.01M phosphate-buffered saline (PBS). Different cellular fraction has been confirmed by Western blot (Supplemental Fig 5c), and protein concentration was measured using Pierce BCA protein Assay Kit (Thermo-Fisher, Mississauga, Ontario, Canada). Samples were stored at −80 °C until use.

### Western blot

Protein expression was assessed using the Wes™ capillary electrophoresis system (Protein Simple, San Jose, CA, USA) according to the manufacturer’s manual. Briefly, samples were diluted to 0.1 μg/μL in provided 1× sample buffer and 1× fluorescent molecular weight marker/reducing agent (Protein Simple). Samples were then vortexed, heat-denatured for 5 min at 95 °C. Samples (0.3 ug) were loaded to each lane into the Wes™ assay plate (Protein Simple), and 12 to 230 kDa separation modules were used. Proteins were separated using capillary electrophoresis and probed with each of the following primary antibodies under default run conditions: anti-GluA1 (1:100; ab109450, Abcam), anti-GluA2 (1:400; ab206293, Abcam), anti-GKAP (1:100; CST #13602. Cell Signaling Technologies, Danvers, MA, USA), anti-Shank3 (1:400; CST #64555, Cell Signaling Technologies), anti-CaMKII (1:400; ab52476, Abcam), anti-phospho-CaMKII(Thr286) (1:400; CST #12716, Cell Signaling Technologies), anti-PSD95 (1:100; CST #3450, Cell Signaling Technologies). Protein expression was normalized to the internal control β-actin (1:100; CST #4970, Cell Signaling Technologies). In order to generate comparable data for all samples, all dilutions of primary antibodies used in Wes™ capillary electrophoresis system have been optimized and validated to ensure it is sufficient to saturate the protein bound to the capillary wall and allow quantitative comparison of signal between samples. Protein was detected using the anti-rabbit detection kit (#DM-001, Protein Simple). Protein was quantified by area under the curve of the chemiluminescent signal. Data was visualized and analyzed using Compass Software version 4.0 for Simple Western (Protein Simple).

### Drugs and Reagents

7-CK was purchased from Abcam (Toronto, ON, Canada). APV, NMDA and capsaicin were purchased from Millipore-Sigma Canada. MTEP, SB-203580, tautomycetin, PYR-41 and UBP-310 were purchased from Tocris Biotechne (Toronto, Ontario, Canada). L689-560 was obtained from Alomone Labs (Jerusalem, Israel). Anisomycin was purchased from Cayman Chemical (Ann Arbor, MI, USA). AV-101 was provided by VistaGen Therapeutics (San Francisco, CA, USA). AV-101 was initially dissolved in 1M NaOH, diluted in phosphate-buffered saline that was titrated to a pH of 7.4 using HCl and injected i.p. at a volume of 10 ml/kg. Cell permeable NMDA receptor mimetics C1.1 corresponding to aa 864-877 of GluN1 (DRKSGRAEPDPKKK), scrambled C1.1 (EPAKDDGRRKPSKK), C1.2 corresponding to aa 877-900 of GluN1 (KATFRAITSTLASS), scrambled C1.2 (AATRSTFASKTLIS) were synthesized with a Tat sequence (YGRKKRRQRRR) at the C-terminal by GenScript (Piscataway, NJ, USA). Scrambled sequences were confirmed to not match other protein sequences in mice by Blast search.

### Statistics

Statistical analyses were performed using GraphPad Prism v9. In all figures, results are expressed as the mean ± standard error of the mean (S.E.M.). All tests were two-sided. MPE was compared between groups using a one way ANOVA followed by Tukey’s multiple comparison post-hoc test or Student’s t-test as appropriate. Withdrawal thresholds in the single capsaicin injection experiment, and CFA hyperalgesia were compared on each day and time point, respectively, using a two-way repeated-measures ANOVA with Bonferroni post-hoc test. Recovery from CFA hyperalgesia for each treatment group was determined by comparing withdrawal thresholds to baseline using one-way repeated-measures ANOVA followed by Dunnett’s test. In biochemical analyses, results were analyzed statistically using one-way analysis of variance (ANOVA) to the vehicle group, followed by a *post hoc* Holm-Sidak test.

Average fPSP areas at baseline, 90 min, and 210 min were compared using a two-way repeated-measures ANOVA with Bonferroni post-hoc test. A D’Agostino-Pearson omnibus normality test was conducted on groups with n > 6, otherwise the Kolmogorov-Smirnov normality test was used. Sample sizes were not predetermined but reflect a balance between sample sizes generally employed in the field and a desire to minimize the use of animals in pain studies. A supplementary methods checklist is available.

## Acknowledgements

This study was funded by grants from the Canadian Institutes of Health Research (CIHR; FRN 162179), the National Science and Engineering Research Council (NSERC) of Canada (RGPIN-2016-05538), the Canadian Pain Society, the University of Toronto Centre for the Study of Pain, and a Canada Research Chair to RPB. We thank Dr. Graham L. Collingridge for his advice on NMDA receptor pharmacology. We thank Maria D’Souza for guidance on the study.

## Supplemental Figures

**Supplemental Figure 1.**
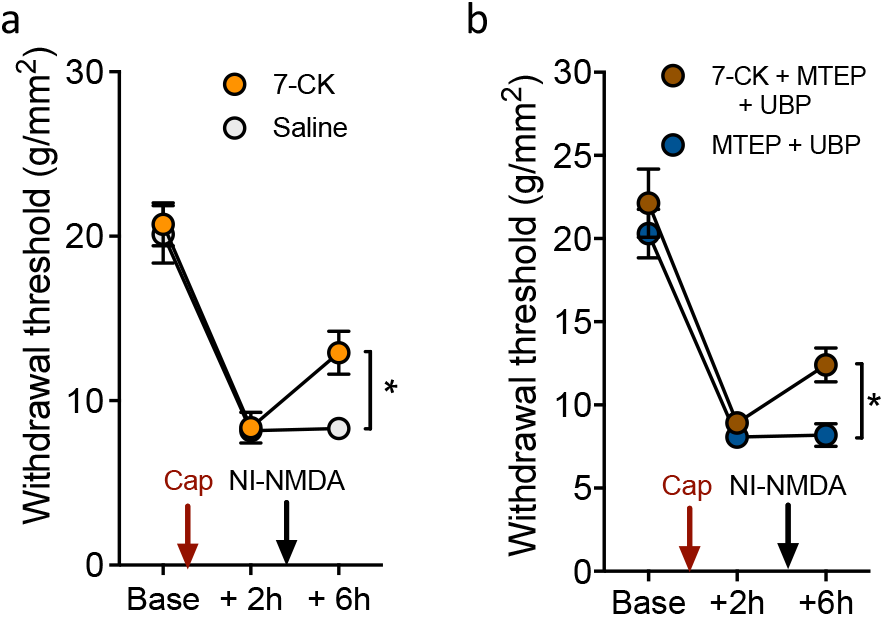
(a) Induction of spinal NI-NMDA with intrathecal 7-CK (250 μM) and plantar capsaicin reverses hyperalgesia in female C57BL/6 mice compared to intrathecal saline and intraplantar capsaicin. n = 7 per group. (b) We examined whether 7-CK reversal of hyperalgesia requires GluK1 kainate receptors or metabotropic glutamate receptor mGluR5. Spinal co-administration of blockers of mGluR5 (MTEP; 200 μM) and GluK1 (UBP-310; 100 μM) with 7-CK did not block the reversal of hyperalgesia after the second plantar capsaicin injection, while these blockers similarly had no effects when spinally administered alone. n = 6 per group, * *p* < 0.05.

**Supplemental Figure 2.**
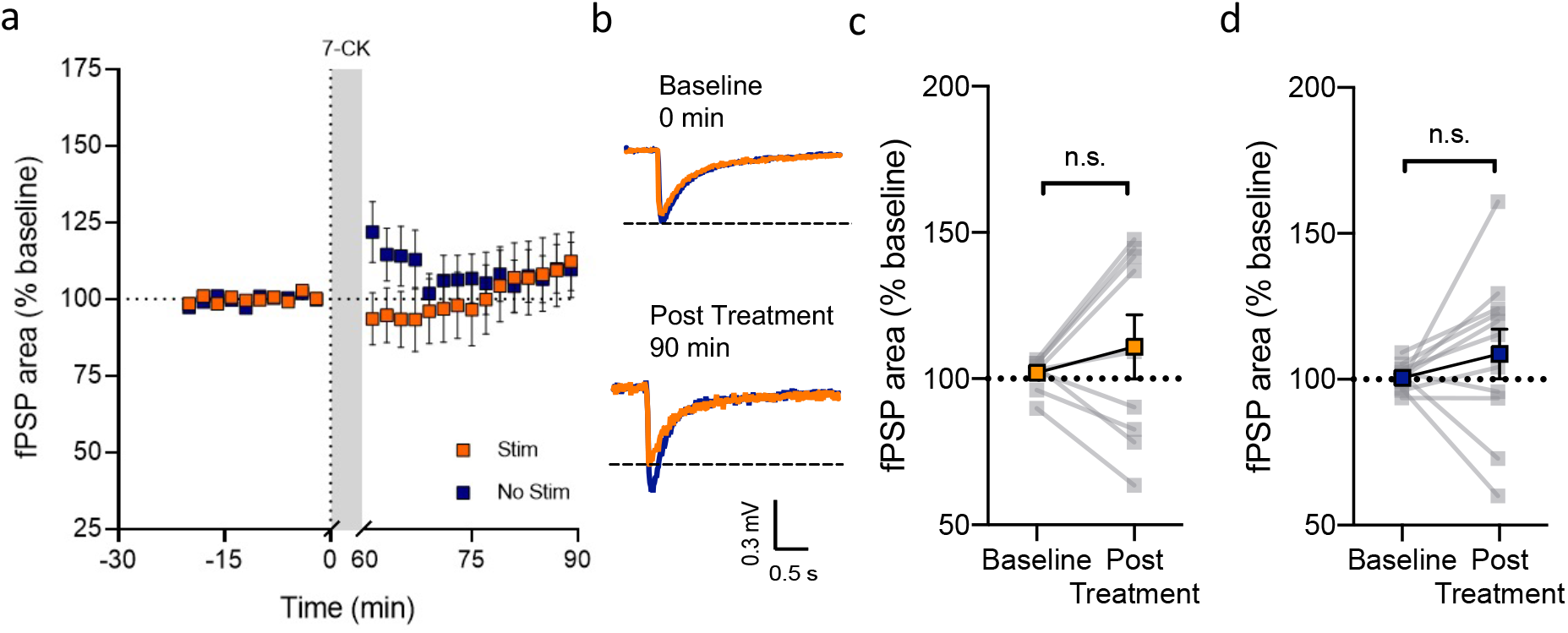
a) After a stable baseline recording, 7-CK was administered from 0 min to 60 min with (orange squares) or without (blue squares) electrical stimulation of dorsal roots. b) Representative traces of fPSPs recorded during baseline (top) and at 90 min (bottom) post 7-CK administration. c-d) fPSP magnitude was compared before (Baseline) and after (Post Treatment) 7-CK administration. Stim: n = 9, t_(8)_ = 0.9089, *p* = 0.3900; No Stim: n = 11, t_(10)_ = 0.9954, *p* = 0.3430.

**Supplemental Figure 3.**
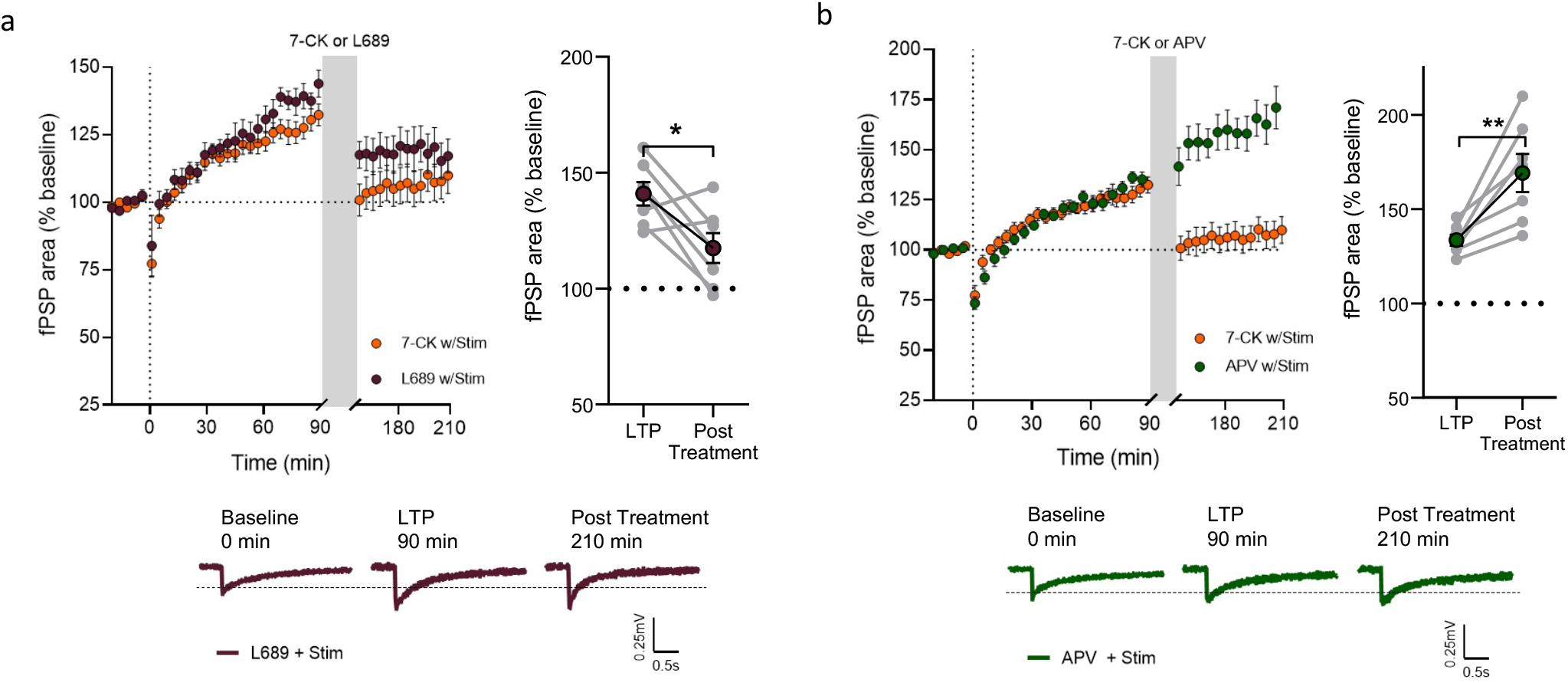
a) Bath application of L689 during stimulation induces activity-dependent depotentiation of dorsal horn LTP in spinal cord explants from female C57BL/6 mice. b) Antagonism at the NMDA receptor glutamate-binding site via APV does not reverse dorsal horn LTP, but instead further potentiates fPSPs after drug administration. 7-CK with stimulation: n = 14, t_(13)_ = 3.514, ** *p* = 0.0038; L689 with stimulation: n = 7, t_(6)_ = 2.845, * *p* = 0.0294; APV with stimulation: n = 7, t_(6)_ = 4.033, ** *p* = 0.0069.

**Supplemental Figure 4.**
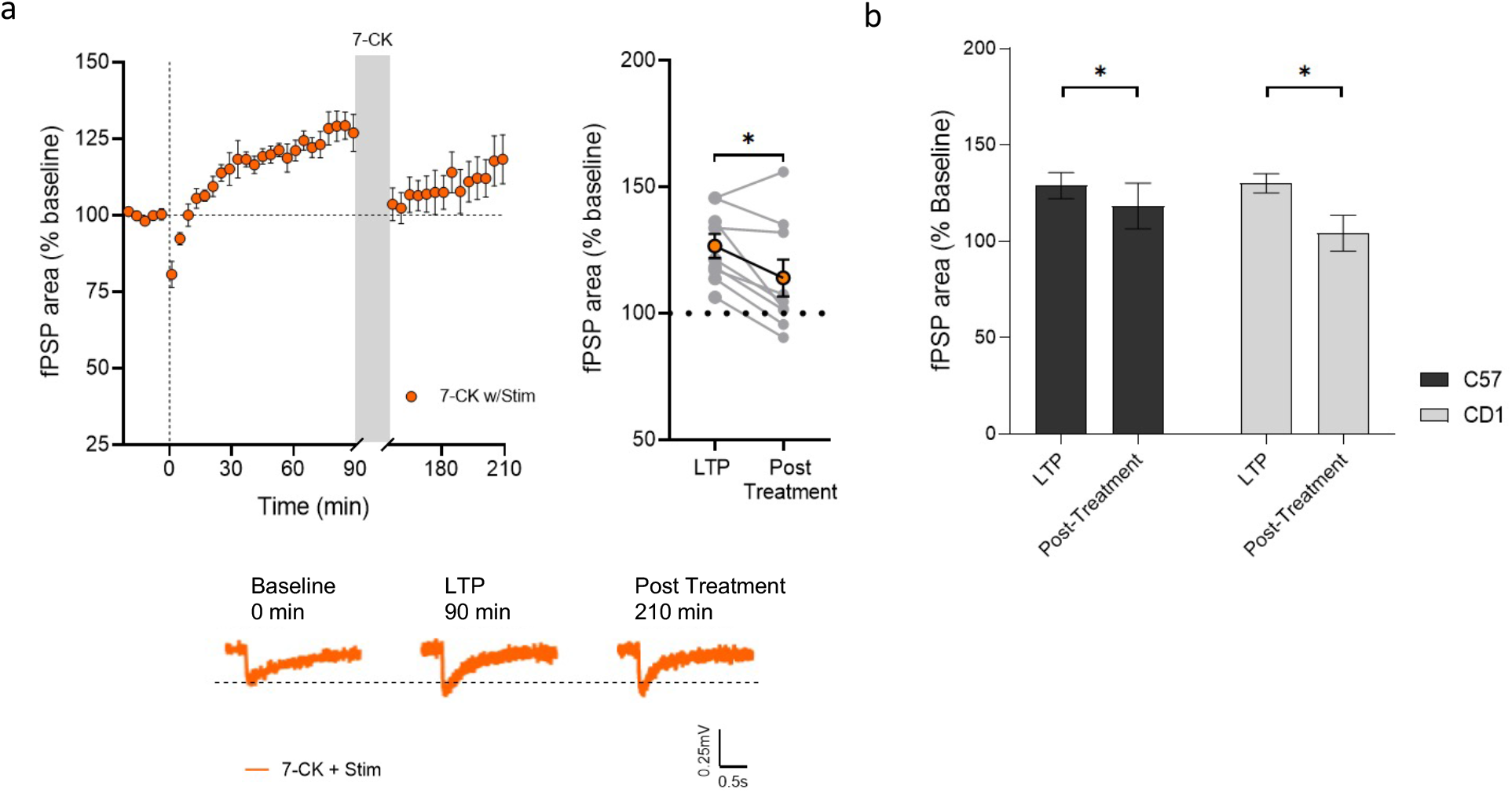
Bath application of 7-CK during stimulation induces activity-dependent depotentiation of dorsal horn LTP in spinal cord explants from female C57BL/6 mice. n = 9, t_(8)_ = 3.077, * *p* = 0.0152. Bath application of 7-CK during stimulation induces a similar degree of depotentiation in spinal explants from C57BL/6 and CD-1 mice. C57BL/6 mice: n = 6, t_(5)_ = 3.078, * *p* = 0.0275; CD-1 mice: n = 7, t_(6)_ = 3.333, * *p* = 0.0158.

**Supplemental Figure 5.**
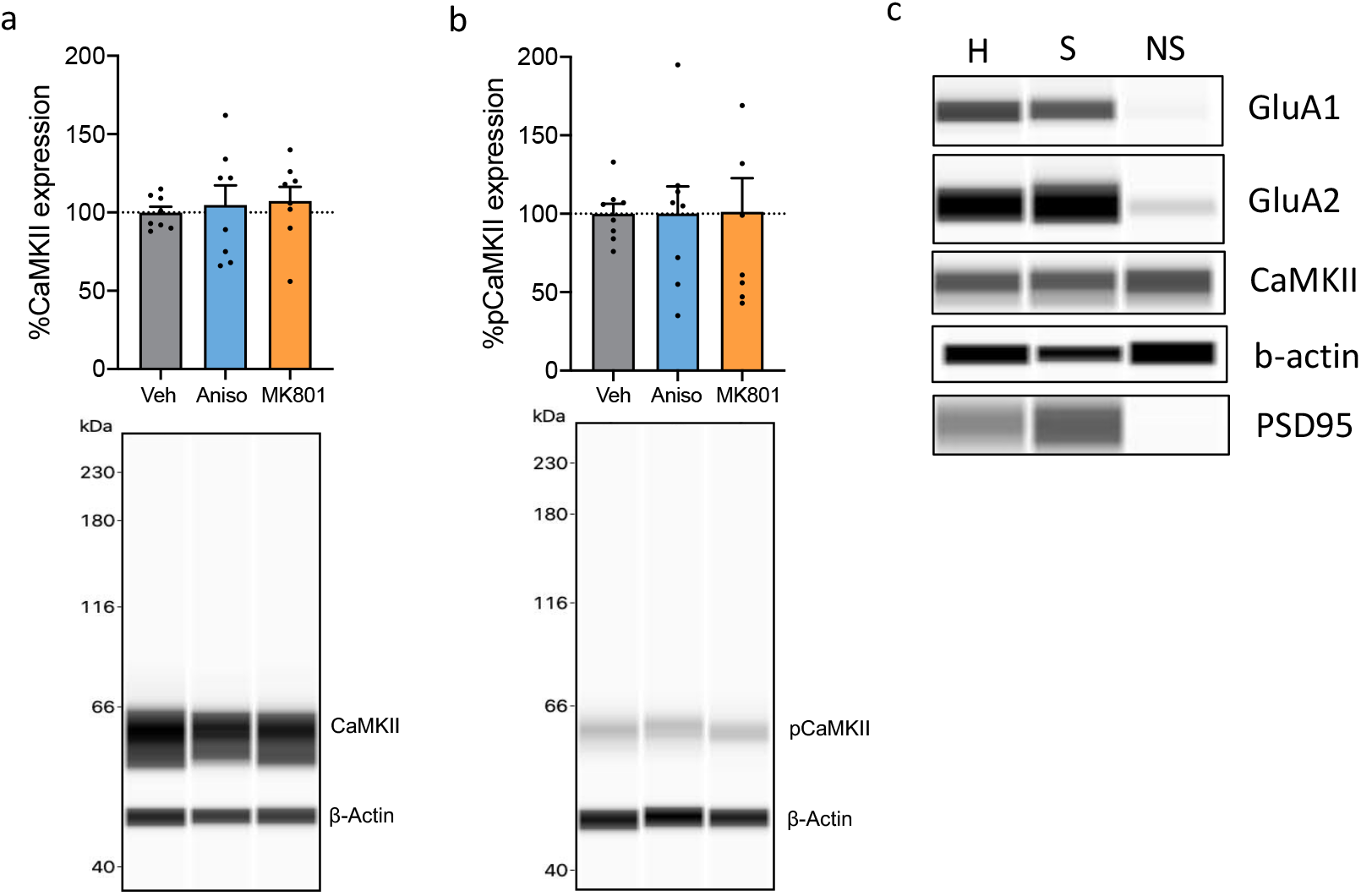
Spinal cord tissue from mice included in Figure 3 were used to generate (a)-(b). The reversal of hyperalgesia by either spinal reconsolidation or NI-NMDA show no change in a) CaMKII (n = 8 per group) and b) Phospho-CaMKII (Thr286) (n = 8 per group) at synaptosome level. c) Synaptic protein expression in different spinal dorsal horn protein fractions; H = homogenized tissue, S = synaptosome fraction, NS = non-synaptic fraction. Western blot analysis of crude synaptosome preparation showed that synaptic markers were strongly maintained in crude synaptosome fraction. Immunoblots of total homogenate (H), crude synaptosome (S), and cytosol non-synaptic fraction (NS) were probed for GluA1, GluA2 (Post-synaptic membrane), CaMKII (synaptic), b-actin (structural, loading control), and PSD95 (post-synaptic density).

